## Supplementary Figures for "Shared and distinct molecular effects of regulatory genetic variants provide insight into mechanisms of distal enhancer-promoter communication"

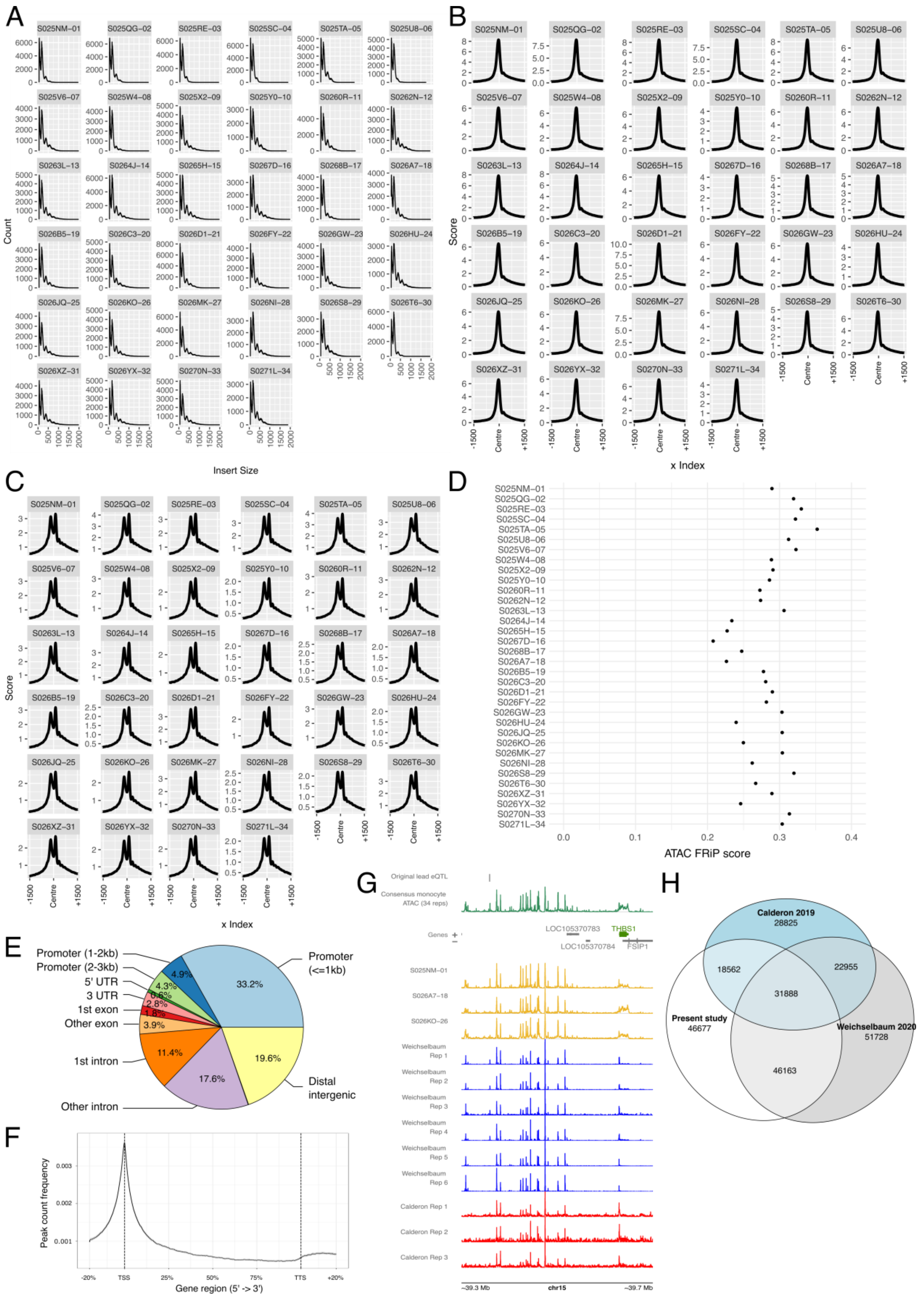


*Supplementary Figure 1.* ***Quality control of ATAC-seq on fixed chromatin in primary monocytes.*** *A) Fragment size plots of ATAC-seq libraries for each replicate. B) Signal profiles of nucleosome-free ATAC-seq reads at transcription start sites for each replicate. C) Signal profiles of mononucleosome ATAC-seq reads at transcription start sites for each replicate. D) Fraction of read in peak (FRiP) score for each*

*ATAC-seq replicate. E) Intersection of ATAC-seq peaks with genomic region annotations (HMMRATAC peaks called using consensus dataset of all 34 replicates). F) Average profile of consensus ATAC peaks binding to genic regions, from 20% upstream of the transcription start site (TSS) to 20% downstream of the transcription termination site (TTS). The 95% confidence interval is shown as a grey ribbon. G) Comparison of ATAC-seq signal in the THBS1 locus: consensus pileup (all 34 replicates), three primary monocyte replicates from the present study and individual replicate-level signal from publicly available primary monocyte ATAC-seq data (Weichselbaum et al., 2020; Calderon et al., 2019). H) Intersecting ATAC-seq peaks between consensus datasets for the present study, Weichselbaum et al. and Calderon et al., determined using Bedtools multiinter on HMRRATAC peaks for all three studies. FRiP, fraction of reads in peak; TSS, transcription start site; TTS, transcription termination site.*

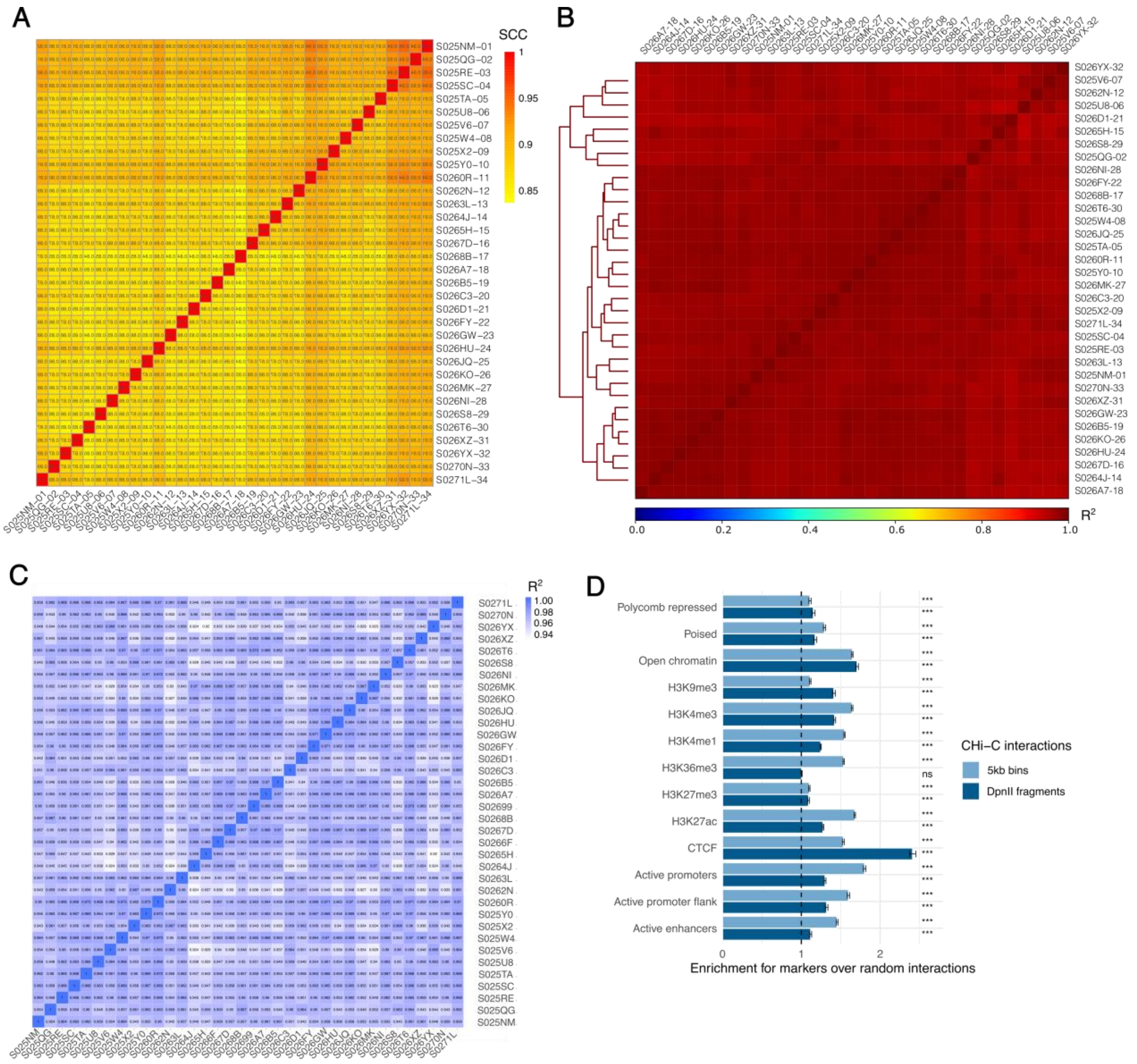


***Supplementary Figure 2. QC of CHi-C, ATAC-seq and RNA-seq samples.*** *A) HiCRep results showing stratum-adjusted correlation coefficient (SCC) values for CHi-C data on chromosome 22. B) Pearson's correlation values for ATAC-seq samples (deepTools). C) Pearson's correlation between RNA samples. Note that sample S02699 was a technical replicate of S026T6. In addition, sample S0266F was a sample that failed genotyping due to a very low call rate, so was not included in any further analyses. D) CHi-C enrichments at other ends of significant interactions at the resolution of 5kb binned regions (light blue) or DpnII fragments (dark blue). For each marker, the enrichment was determined against randomly sampled chromatin fragments located the same distance away from the baited regions as the detected interacting fragments, using the peakEnrichment4Features function in Chicago[33,34]. *** indicates p-values < 0.001 and "ns" represents "non significant" (permutation test with 100 random samples). Error bars show the standard deviation of the mean ratio between true interacting fragments and the random interacting fragments. Histone modification peaks were obtained from Blueprint. Further annotations (promoter flanking regions, promoters, enhancers, poised and polycomb repressed regions) were taken from Blueprint projected segmentations in monocytes. ATAC-seq peaks of open chromatin were obtained from the present monocyte cohort (consensus HMMRATAC peaks).*

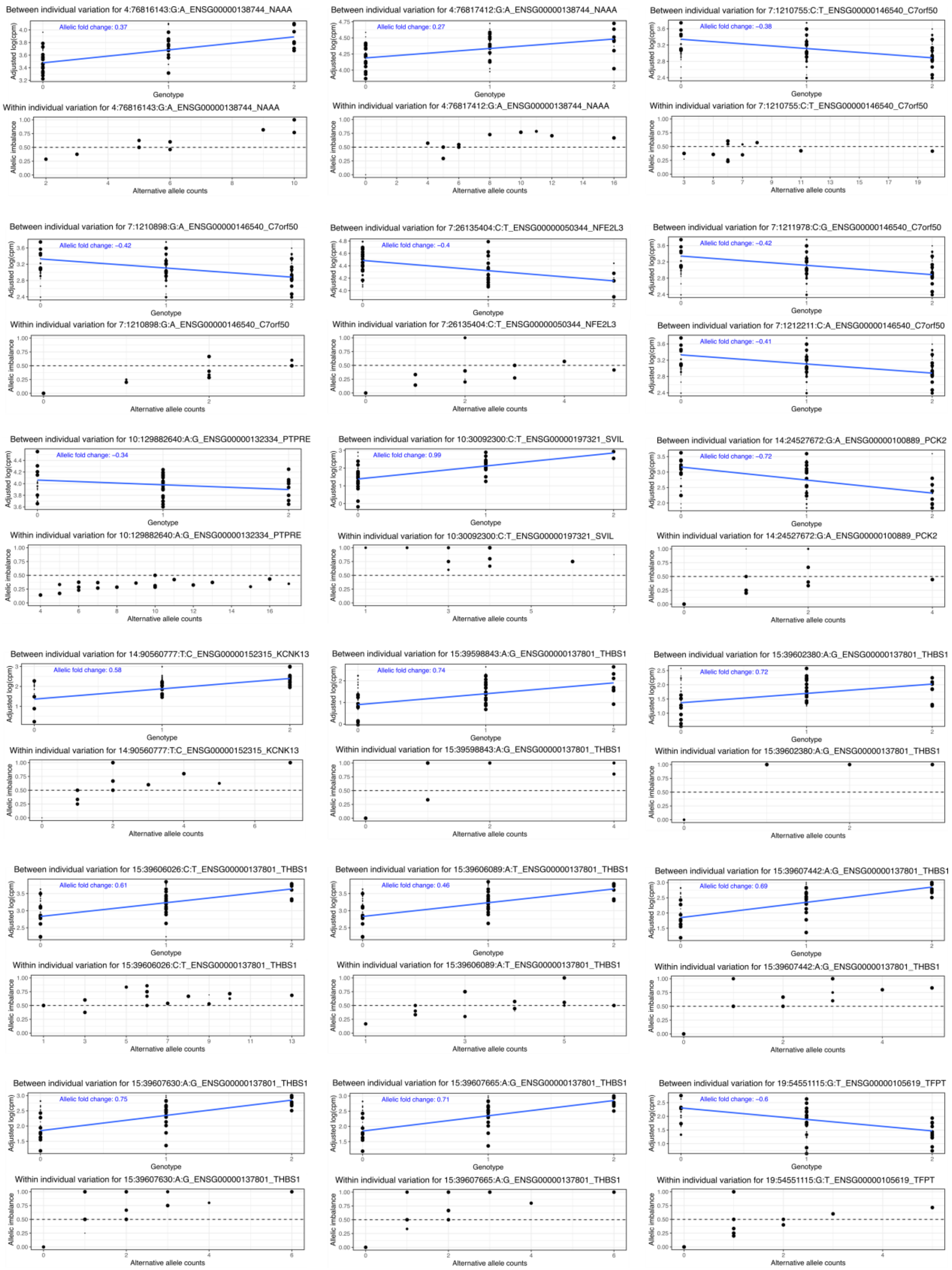


***Supplementary Figure 3. BaseQTL estimated effects for CHi-C QTLs.*** *For each of the 19 significant BaseQTL contact eQTLs, we illustrate the two components of our model: between-individual (top) and within-individual (bottom) variation. Between-individual plots show the genotype of the cis-SNP (x-axis) against the total counts adjusted by library size in log scale (y-axis). The size of each point corresponds to the probability of the indicated genotype. To represent the within-individual variation, only heterozygotes are considered. The x-axis corresponds to the number of reads mapping the cis-SNP alternative allele, while the y-axis shows the proportion of reads mapping the cis-SNP alternative*

*allele. The size of each point corresponds to the probability of the cis-SNP genotype being heterozygous. The value of the estimated log allelic fold change is given in blue. For the associations between cis-SNPs 7:1211978:C:G and 7:1212211:C:A with the promoter of C7orf50, only between-individual information was available to model.*

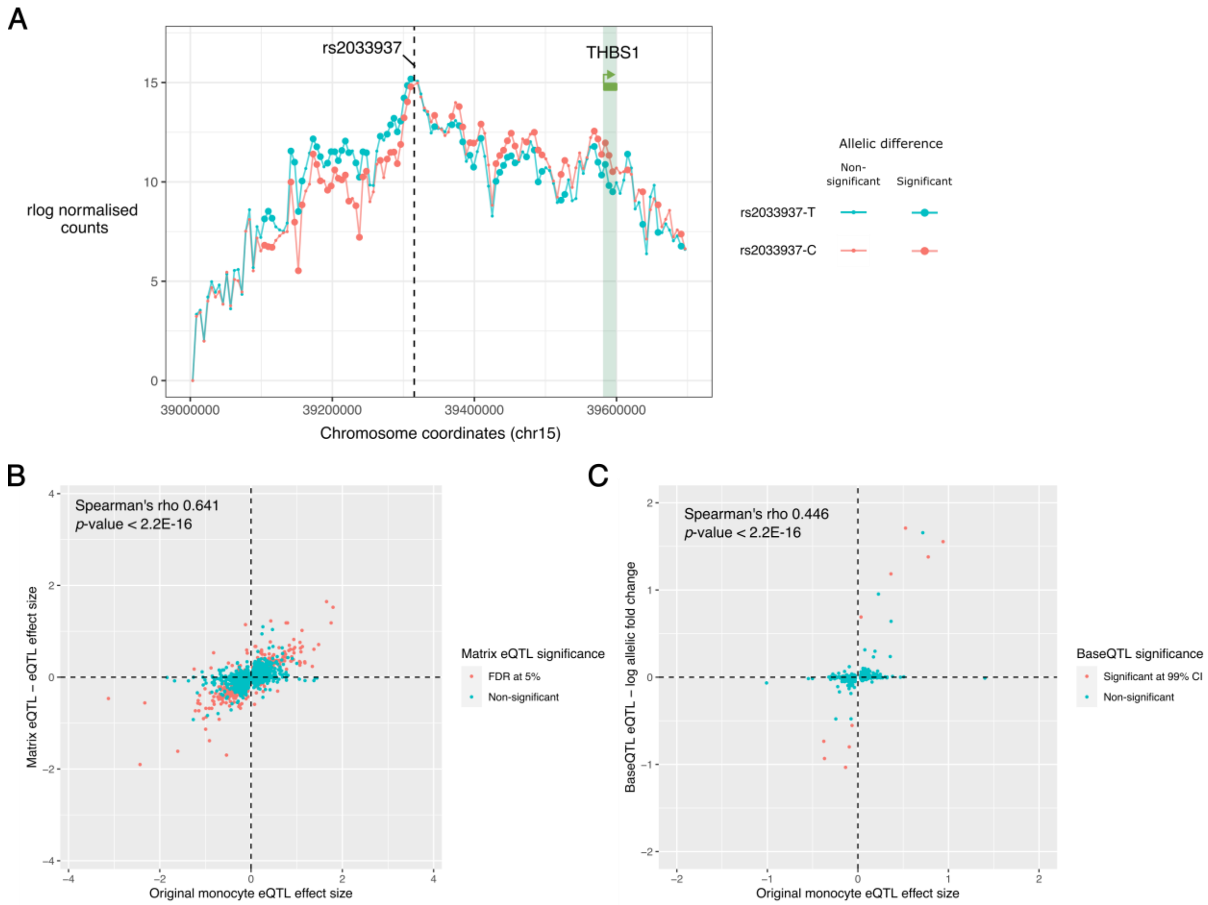


***Supplementary Figure 4. Validation of a chromatin QTL at THBS1 and correlation between eQTL effects in the original multi-cohort monocyte study and in the present cohort**. A) 4C-seq validation of allele-specific chromatin conformation between the eQTL rs2033937 and its eGene THBS1. The mean 4C-seq signal across three heterozygous individuals (rlog normalised reads) is shown for the reference allele (T, blue) and the alternative allele (C, red) of rs2033937. Significantly different interactions are shown as large dots (4Cker adjusted p-value $< 0.05$). B) Correlation of the original eQTL dataset with eQTLs called by Matrix eQTL in the present study. C) Correlation of the original eQTL dataset with eQTLs called by BaseQTL in the present study.*

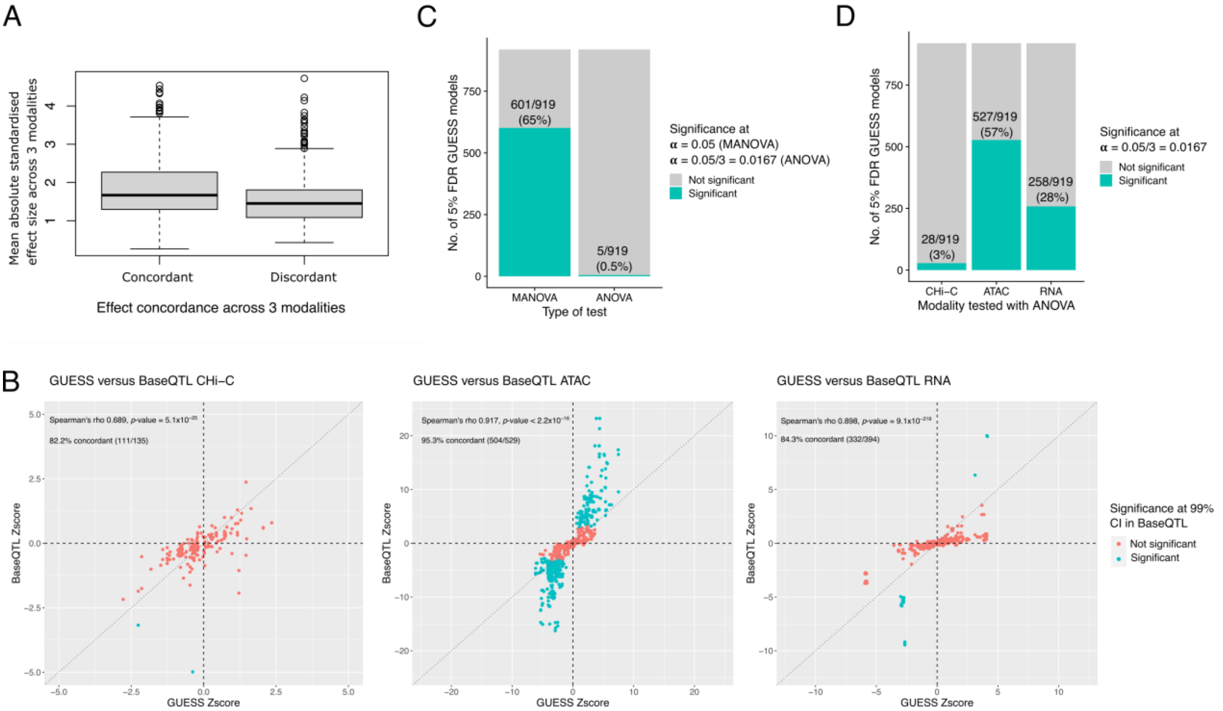


***Supplementary Figure 5. Validation of GUESS results.*** *A). Standardised effect size of GUESS models depending on overall concordance. In concordant models, the sign of the beta was the same for all three modalities (CHi-C, ATAC-seq and RNA-seq). For discordant models, at least one of the signs was different. Only GUESS models containing one trimodal QTL at 5% FDR were included in this analysis. B) GUESS and BaseQTL concordance. The GUESS data (FDR 5%, single QTL models) and BaseQTL data (all results) were merged on the trimodal QTL and the feature: CHi-C bait fragment (left), ATAC-seq peak (middle) or gene (right). The results are coloured by their significance in BaseQTL at a CI cutoff of 0.99. Only single-QTL models in GUESS were used for the comparisons in this figure. Concordance is defined as a QTL having the same direction of effect across analyses. C) We performed multivariate analysis of variance (MANOVA) on the lead trimodal QTL identified by GUESS in each window, treating the other trimodal QTLs in the same window (if detected at 5% FDR) as covariates (MANCOVA), with the graph showing the number of models at alpha = 0.05. We also assessed how many of the lead trimodal QTLs could be validated using a univariate approach (ANOVA/ANCOVA), with the graph showing the number of models at alpha = 0.0167 for all three modalities (adjusting for multiple testing). D) Validation of each of the three individual modalities using ANOVA/ANCOVA (alpha = 0.0167).*

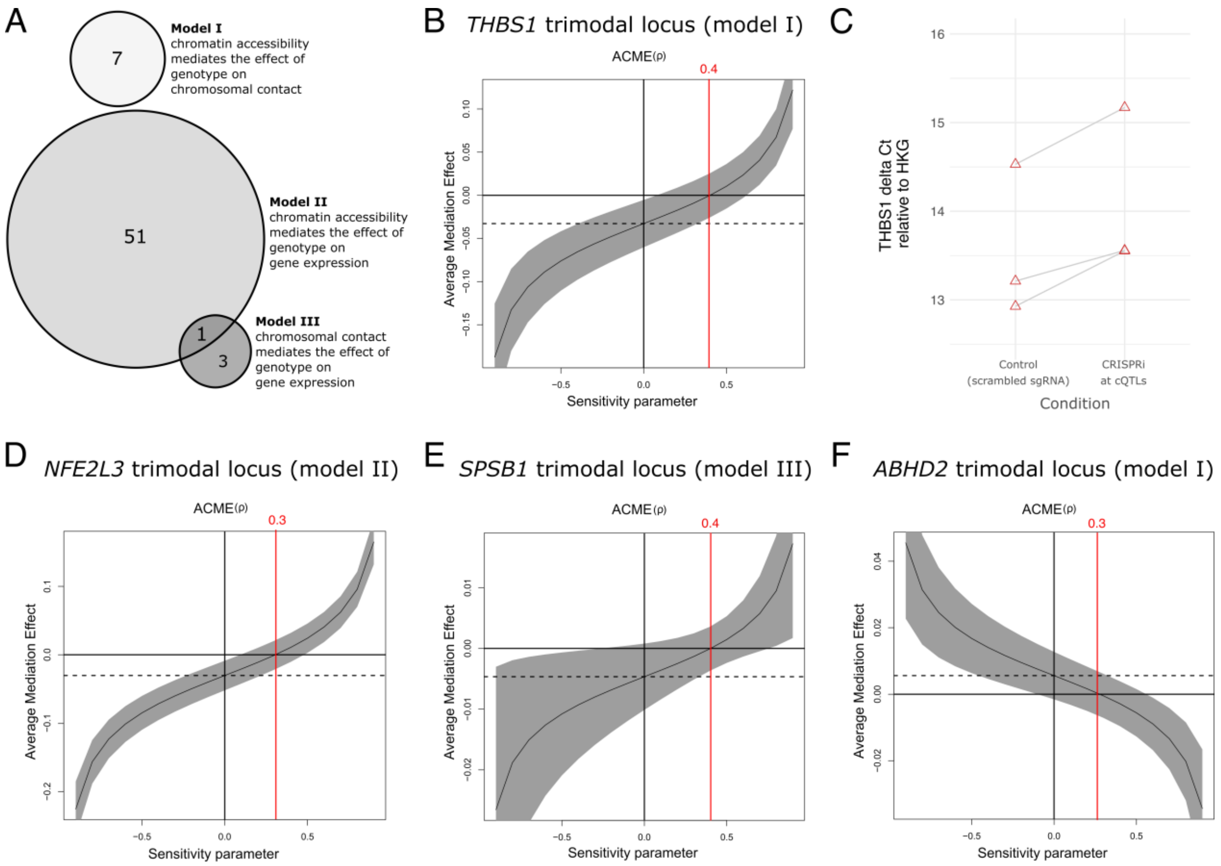


***Supplementary Figure 6. Mediation models.*** *A) Total Number of trimodal QTLs implicated in each mediation model. B). Sensitivity plot testing for the existence of unobserved post-treatment confounders in the THBS1 locus. The plot shows the values of causal quantities as a function of sensitivity parameter (the correlation between the residuals of the mediator and outcome regressions). The solid line in the curve represents the estimated average mediation effect at different values of the sensitivity parameter, with the grey areas representing the 95% confidence interval. The ACME estimate is shown as a black horizontal dotted line. The red vertical line shows the level of confounding required to observe no mediation effect (ACME = 0) and, thus, invalidate the results. Values ≥ 0.3 indicate that a strong confounding effect is necessary to change the sign of ACME estimates. C) Delta Ct values for the THSB1 expression qPCR in the CRISPRi experiment, compared with the housekeeping genes (HKG) GAPDH, TOP1 and ATP. D-F) same as B but for the NFE2L3, SPSB1 and ABDH2 loci.*

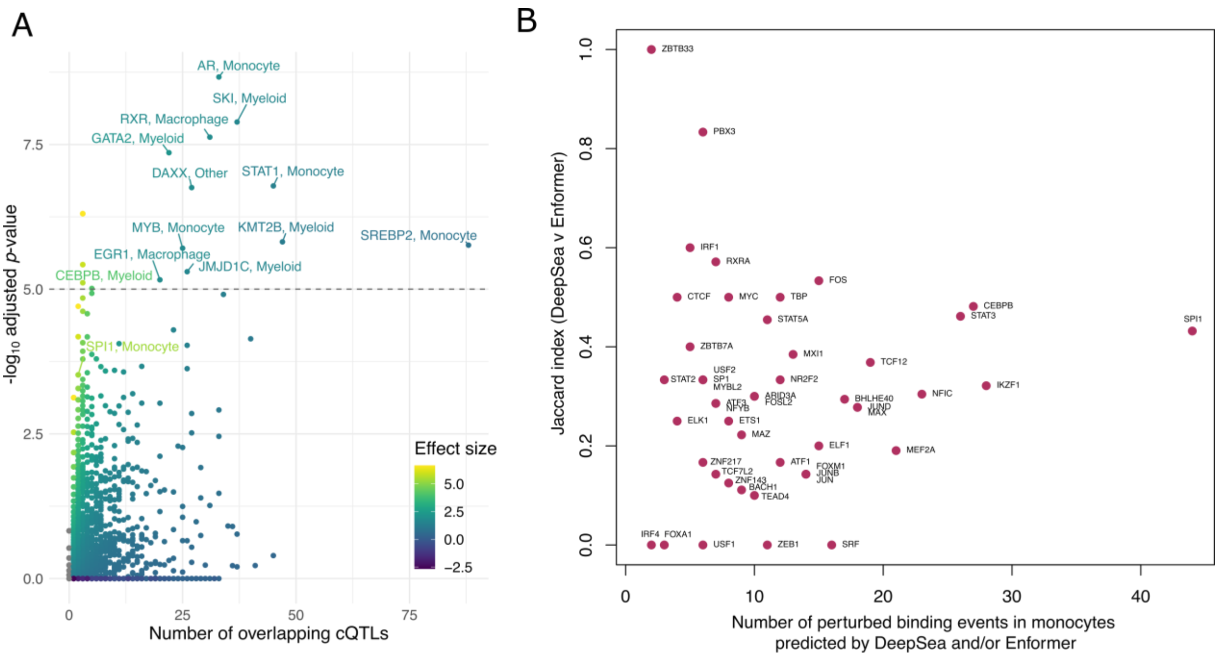


***Supplementary Figure 7. cQTLs intersect and are predicted to perturb TF binding sites****. A) Enrichment of experimentally determined TF binding sites at cQTLs (ReMap 2022 ChIP-seq dataset). TFs are labelled at -log10 adjusted p-value (Q significance) > 5 and number of overlapping cQTLs >= 5, apart from SPI1, which is shown for reference, because it was also identified in the analysis of monocyte-specific ChIP-seq and ATAC-seq predicted peaks. Each cell type was classified as "monocyte", "macrophage", "myeloid" or "other". B) Jaccard index of DeepSea vs Enformer is shown against the number of predicted perturbed binding events in monocytes at cQTLs, as predicted by DeepSea and/or Enformer.*
